# Yersiniabactin, Colibactin and Wider Resistome Contribute to Enhanced Virulence and Persistence of KPC-2-Producing *Klebsiella pneumoniae* CG258 in South America

**DOI:** 10.1101/435750

**Authors:** Louise T. Cerdeira, Fernanda Esposito, Margaret M. C. Lam, Kelly L. Wyres, Ryan R. Wick, Louise M. Judd, Marcos V. Cunha, Terezinha Knöbl, Bruna F. Araújo, Rosineide M. Ribas, Sebastian Cifuentes, Jesus Tamariz, Maria F. C. Bueno, Gabriela R. Francisco, Doroti O. Garcia, Gerardo Gonzalez-Rocha, Gabriel Gutkind, Kathryn E. Holt, Nilton Lincopan

## Abstract

The emergence and dissemination of carbapenem-resistant hypervirulent *Klebsiella pneumoniae* (CR-hvKp) is a worrisome public health issue compromising the treatment and outcome of infections caused by this pathogen. We performed a detailed virulome and resistome analysis of representative KPC- and/or CTX-M-producing *K. pneumoniae* belonging to clonal group (CG) 258 (sequence types ST11, ST258, ST340, ST437), circulating in Argentina, Brazil, Chile, Colombia and Peru; with further evaluation of the virulence behavior using the *Galleria mellonella* infection model. Genomic analysis of *K. pneumoniae* strains recovered from the human-animal-environment interface revealed a wide resistome characterized by the presence of genes and mutations conferring resistance to human and veterinary antibiotics, quaternary ammonium compounds (QACs) and heavy metals. Plasmid Inc typing revealed the presence of a wide diversity of replicon types with IncF, IncN, IncR and Col-like being frequently detected. Moreover, KPC-2-producing *K. pneumoniae* belonging to ST11 (KL-64 andKL-105) and ST340 (KL-15) carried multiple variants of distinct yersiniabactin siderophore (*ybt*) and/or genotoxic colibactin (*clb*) genes. In this regard, ICE*Kp3*, ICE*Kp4* and ICE*Kp12* were identified in strains belonging to ST11 and ST340, recovered from Argentina, Brazil, Chile and Colombia; whereas *ybt* 17 and a novel *ybt* sequence type (YbST346) were identified together with *clb* in ICE*Kp10* structures from ST11 and ST258, from Brazil and Colombia, respectively. *K. pneumoniae* ST11 (ICE*Kp10*/YbST346 and ICE*Kp4*/*ybt* 10) strains killed 100% of wax moth larvae, in a similar way to hypervirulent K1/ST23 strain (*ybt*- and *clb*-negative) carrying the pLVPK-like plasmid, indicating enhanced virulence. In summary, our results indicate that yersiniabactin, colibactin and an expanded resistome have contributed to enhanced virulence and persistence of KPC-2-producing *K. pneumoniae* CG258 in South America. Therefore, active surveillance of hospital-associated lineages of *K. pneumoniae* should not only focus on clonal origin and antimicrobial resistance, but also on the virulence factors *ybt* and *clb*.

## INTRODUCTION

Carbapenem resistance is a major public health concern worldwide, and currently *Klebsiella pneumoniae* belonging to the clonal group CG258 (which include the sequence types ST11, ST258, ST340, ST437, and ST512) seem to be the main culprits for the spread of bla_KPC_ genes (Bowers et al., 2015; Chen et al., 2014; Holt et al. 2015; Mathers et al., 2015; Paczosa and Mecsas 2016; Wyres and Holt, 2016). This problem has been further exacerbated by the convergence of KPC-2 production and hypervirulence, resulting in the emergence of carbapenem-resistant hypervirulent *K. pneumoniae* (CR-hvKp) lineages, particularly in Asian countries (Chen et al., 2017; Lee et al., 2017; Dong et al., 2018a; 2018b; Du et al., 2018; Gu et al., 2018; Wang et al., 2018). In these countries hypervirulence has been associated with the appearance and dissemination of a pLVPK-like plasmid harbouring two capsular polysaccharides (CPS) upregulator genes (*rmpA* and *rmpA2*) and several siderophore gene clusters (*iroBCDN*, *iucABCD* and *iutA*) (Struve et al., 2015; Chen et al., 2017; Du et al., 2018; Gu et al., 2018). However, the acquisition of integrative conjugative elements (ICE*Kp*) harbouring yersiniabactin siderophore (*ybt*) is also associated with enhanced virulence, whereas carriage of the genotoxic colibactin (*clb*) genes (in ICE*Kp10*structures) has been associated with invasive disease and colorectal cancer (Holt et al., 2015; Lam et al., 2018a).

Based on gene content variation, genomic investigation has allowed the identification of 14 different structural ICE*Kp* variants, constituting a novel target that deserves further analysis for evolutionary and genomic surveillance studies (Wu et al., 2009; Lam et al., 2018a; Lin et al, 2008). A MLST-style approach based on diversity in eleven *ybt* locus genes has defined yersiniabactin sequence types (YbSTs) by unique combinations of *ybt* gene alleles and showed that YbST sequences were clustered into 17 distinct *ybt* lineages (Lam et al., 2018a). In a similar way, variations in the *clb* locus genes have allowed definition of colibactin sequence types (CbSTs), whereas phylogenetic analysis of the *clb* locus has revealed three lineages that have each associated with a different *ybt* lineage [i.e., *clb* 1 (*ybt* 12), *clb* 2A (*ybt* 1) and clb 2B (*ybt* 17)] within the same overall structure (ICE*Kp10*).

Virulence in CR-Kp strains has also been associated with the type of capsular polysaccharide (Cortés et al., 2002; Diago-Navarro et al., 2014; Gomez-Simmonds and Uhlemann, 2017; Liu et al., 2017). In this regard, over 79 capsule (K) serotypes have been described in the international K serotyping scheme (Brisse et al., 2004; Pan et al., 2015; Struve et al., 2015). More recently, diversity of the capsule synthesis locus (K-locus), which is 10–30 kbp in size, has been used as a novel typing method for genomic surveillance and epidemiological investigations of this pathogen, and identified 134 distinct K-loci, which are predictive of K serotype (Wyres et al., 2016).

In South American countries, KPC-2-producing *K. pneumoniae* has been circulating in Colombia, Brazil and Argentina since at least 2005 (Villegas et al., 2006; Pavez et al., 2009; Gomez et al., 2011), and has more recently been reported in Ecuador, Chile, Venezuela, Paraguay, Uruguay and Peru (Cifuentes et al., 2012; Zurita et al., 2013; Marquez et al., 2014; Falco et al., 2016; Gomez et al., 2016; Horna et al., 2017). Although, molecular epidemiology studies have confirmed predominance of the CG258 among KPC-2-producing *K. pneumoniae* isolates collected in this region (Andrade et al., 2011; Cejas et al., 2012; Pereira et al., 2013; Gomez et al., 2016; Barría-Loaiza et al., 2017; Horna et al., 2017), few studies have focused on virulence determinants in these strains (Andrade et al., 2018; Araújo et al., 2018). In fact, studies of biofilm formation and identification of a common set of virulence genes have been restricted to KPC-2-producing *K. pneumoniae* from Brazil, whereas sporadic identification of hypervirulent *K. pneumoniae* (hvKp) isolates belonging to K1/ST23 and K19/ST29 have been reported in infected patients from Argentina and Brazil (Cejas et al., 2014; Coutinho et al., 2014; Moura et al., 2017), and also in non-human primates from Brazil (Anzai et al., 2017). In this study, using a genomic approach, we have performed a detailed virulome and resistome analysis of KPC- and CTX-M-producing *K. pneumoniae* strains belonging to CG258, recovered from the human-animal-environment interface in Latin America, with further *in vivo* virulence evaluation using a *Galleria mellonella* infection model.

## METHODS

### *K. pneumoniae* strains and genome collection

Laboratory studies included 19 KPC-2- and/or CTX-M-producing *K. pneumoniae* isolates belonging to CG258 (ST11, ST258, ST340, ST437), representative of local surveillance studies performed in Brazil, Peru, Chile and Argentina, between 2010 to 2016; recovered from human, food-producing animals (chicken and swine) and environmental (urban rivers and urban lake) samples (Oliveira et al., 2014; Martins et al., 2015; Cerdeira et al., 2016a;2016b; Cerdeira et al., 2017; Horna et al., 2017; Nascimento et al., 2017). *K. pneumoniae* ATCC 13883 and hvKp K1/ST23 A58300 (Coutinho et al., 2014) were used as control strains. For genome analysis, all publicly available genomes from 36 *K. pneumoniae* CG258 strains isolated in South America were included, of which 23 were previously published (Bowers et al., 2015; Araújo et al., 2018; Casella et al., 2018; Dalmolin et al., 2018; Pitt et al., 2018). For all 55 genomes included in this work, accession numbers are listed in Table S1.

#### Antibiotic susceptibility patterns and hypermucoviscosity phenotypical identification

Resistance phenotypes were determined by Kirby-Bauer method, against 30 different human and veterinary antibiotics, and the results were interpreted using the Clinical and Laboratory Standards Institute guidelines (CLSI, 2015; 2017) and The European Committee on Antimicrobial Susceptibility Testing (EUCAST, 2017). Additionally, minimum inhibitory concentrations (MICs) for ertapenem, imipenem, meropenem, enrofloxacin, ciprofloxacin, levofloxacin and polymyxin B, were determined by microdilution or Etest methods (CLSI, 2017; EUCAST, 2017). Production of ESBL and carbapenemase enzymes was confirmed by growth on CHROMagar ESBL and CHROMagar KPC, respectively.

Hypermucoviscosity phenotypes of *K. pneumoniae* isolates were determined by the string test as previously described (Moura et al., 2017). Briefly, a positive string test was defined by the formation of a viscous string > 5 mm in length when a colony was grown on a blood agar plate at 37 °C overnight and stretched by an inoculation loop.

### Sequencing of *K. pneumoniae* strains

For 12 *K. pneumoniae* strains, total genomic DNA was extracted using PureLink™ Genomic DNA Mini Kit according to the Manufacturer’s instructions. Library preparation was performed using the Nextera XT DNA sample preparation kit (Illumina, San Diego, CA). Sequencing was performed using the Illumina NextSeq platform with paired-end reads (150bp). Additionally, 10 *K. pneumoniae* isolates selected from the 19 KPC-2 and/or CTX-M producers (Table S1) were subjected to further sequencing using a PromethION R9.4.1 flow cell (Oxford Nanopore Technologies). A 2D MinION library was generated from 1.5 µg purified genomic DNA using the Nanopore Sequencing Kit (SQK-NSK007). DNA was repaired (NEBNext FFPE RepairMix), prepared for ligation (NEBNextUltra II End-Repair/dA-tailing Module) and ligated with adapters (NEB Blunt/TA Ligase Master Mix).

### Bioinformatic analysis

Forty-one *K. pneumoniae* CG258 genomes with available short-read sequence data (including genomes obtained in this study and publicly available in the GenBank) were subjected for *de novo* assemblies using Unicycler (v0.4.0) (Wick et al., 2017). For ten genomes obtained with Nanopore reads, scaffold bridging was performed, building a high-quality finished genome sequence. The contigs were annotated by Prokka v1.12 (https://github.com/tseemann/prokka). MLSTs, YbSTs, CbSTs (Lam et al., 2018a; Diancourt et al., 2005), virulome, resistome and plasmid replicon genes were screened by SRST2 (Inouye et al., 2014), using BIGSdb (Bialek-Davenet et al., 2014), ARG-Annot (Gupta et al., 2014), and PlasmidFinder (Carattoli et al., 2014) databases. On the other hand, since 14 of the publicly available genomes (used in this study) were only available as pre-assembled sequences, Kleborate (https://github.com/katholt/Kleborate) and PlasmidFinder were used to identify MLST, resistome, virulome and plasmid replicon genes. Kleborate was further used to predict yersiniabactin ICE*Kp* structures in all 55 genomes.

ResFinder 3.0 database was used to confirm resistomes (Zankari et al., 2012), and while capsule and O-antigen biosynthesis loci were identified using Kaptive (Wick et al., 2018), heavy metal (HM) and QAC genes were screened using BLASTN against local HM/QAC and BIGSdb databases.

Single nucleotide variants were identified using RedDog v1beta.10.3 (https://github.com/katholt/RedDog), with the reference genome of *K. pneumoniae*30660/NJST258_1 (CP006923) (Bowers et al., 2015). Gubbins v.2.1.0 (Croucher et al., 2014) was used to identify and exclude recombination imports. Maximum likelihood (ML) trees were inferred from the recombination-masked alignment by running RaxML v8.2.9 (Stamatakis et al., 2006) five times, selecting the final tree with the highest likelihood. To assess branch support, we conducted 100 non-parametric bootstrap replicates using RAxML.

### *Galleria mellonella* killing assays

In order to evaluate the virulence behavior of KPC-2- and/or CTX-M-15-producing *K. pneumoniae* strains, *in vivo* experiments were carried out using a Galleria mellonella infection model (Junqueira 2012; Insua et al. 2013), with the non-virulent *K. pneumoniae* strain ATCC 13883 and the clinical hvKp K1/ST23 strain A58300 (Coutinho et al. 2014), as comparative strains. Fourteen *K. pneumoniae* strains of different lineages of CG258 and origins (i.e., human, animal or environmental), circulating in Latin America, were evaluated. For each experiment, a control group containing five larvae was inoculated with sterile PBS in order to discard death due to physical trauma. In all experiments, groups of 250 to 350 mg G. mellonella larvae were inoculated with 10^6^ CFU, and survival analysis was evaluated during 96h (Moura et al. 2017). Survival curves were plotted using the Kaplan-Meier method, and data were analyzed by the Fisher’s exact test, with P< 0.001 indicating statistical significance. The statistical software used was Prism7 (Graph Pad Software, San Diego, CA, USA). *G. mellonella* larvae that did not demonstrate a response to physical stimulation and had body melanization were considered dead. All experiments were performed in independent triplicate assays.

## RESULTS

### Antimicrobial resistance profiles, resistome and plasmid populations

All 19 *K. pneumoniae* evaluated, *in vitro*, exhibited resistance to multiple antibiotics and were classified as MDR or PDR phenotypes (Magiorakos et al., 2012) (Table 1). In fact, resistome analysis revealed the presence of genes conferring resistance to aminoglycosides, quinolones, sulphonamides, tetracycline, phenicols, fosfomycin and beta-lactam antibiotics (*bla*_OXA-1_, *bla*_OXA-2_, *bla*_OXA-9_, *bla*_OXA-10_, *bla*_CTX-M-15_, *bla*_SHV-11_, *bla*_SHV-12_, *bla*_LAP-2_, *bla*_TEM-1A_, *bla*_TEM-1B_, *bla*_TEM-55_) (Figure 1; Table S2). Moreover, point mutations in GyrA (Ser-83-Ile), GyrB (Asp-466-Glu) and ParC (Ser-80-Ile) were associated with quinolone resistance. Polymyxin resistance in 5 of the 19 (26%) human and environmental isolates was associated to *mgrB* mutations (i.e., Gly-28-Cys or Tn3 insertion at position 134) (Table S2). Additionally, the presence of genes conferring resistance to silver (*sil*), copper (*pco*), arsenic (*ars*), mercury (*mer*), tellurite (*ter*), and quaternary ammonium compounds (*qacA*, *qacE*, *qacE∆1*, *qacL* and *sugE*) supported a wider resistome (Figure 1, Table S2), which could contribute to the apparent high versatility, persistence and adaptation of CG258 to various ecosystems and hosts (Navon-Venezia et al., 2017; Dong et al., 2018). Notably, the presence of tellurite resistance has also been associated with hypervirulent clonal groups of *K. pneumoniae* (Passet and Brisse, 2015; Martin et al., 2018).

**Table 1.** Resistance profile of KPC-2- and/or CTX-M-producing *K. pneumoniae* strains belonging to CG258 circulating in South America*

| Strain | MICs (µg/ml) |  |  |  |  |  |  | Kirby-Bauer |
| --- | --- | --- | --- | --- | --- | --- | --- | --- |
|  | POL | ETP | IMP | MER | ENO | CIP | LVX |  |
| KP171 | 2 | 8 | 16 | 8 | 8 | 8 | 32 | CTX, CAZ, CPM, CRO, NAL, GEN, SUT, ATM, TET, CLO, FOS |
| 1194 | 2 | 4 | 8 | 8 | 8 | 8 | 32 | CTX, CAZ, CPM, CRO, NAL, AMI, GEN, SUT, ATM, TET, CLO, FOS |
| 606B | 2 | 1 | 1 | 0.25 | 4 | 4 | 16 | CTX, CAZ, CPM, CRO, NAL, GEN, SUT, ATM, TET, CLO, FOS |
| KPN535 | 16 | 4 | 8 | 4 | 8 | 8 | 32 | CTX, CAZ, CPM, CRO, NAL, GEN, SUT, ATM, CLO, FOS |
| KPC45 | 32 | 4 | 8 | 4 | 4 | 4 | 16 | CTX, CAZ, CPM, CRO, NAL, SUT, ATM |
| IBL2.4 | 8 | 4 | 8 | 8 | 4 | 8 | 32 | CTX, CAZ, CPM, CRO, CFO, NAL, ATM, TET |
| KP488 | >32 | 4 | 8 | 8 | 4 | 4 | 32 | CTX, CAZ, CPM, CRO, CFO, NAL, GEN, SUT, ATM, CLO, FOS |
| 196 | 0.25 | >32 | 8 | >32 | 8 | >32 | >32 | CTX, CAZ, CPM, CRO, CFO, NAL, SUT, ATM |
| 148 | 32 | >32 | >32 | >32 | 4 | >32 | >32 | CTX, CAZ, CPM, CRO, CFO, NAL, SUT, ATM, TET, CLO, FOS |
| 314 | 2 | >32 | >32 | >32 | 4 | >32 | >32 | CTX, CAZ, CPM, CRO, CFO, NAL, SUT, ATM, TET |
| KP411 | 0,5 | >32 | >32 | >32 | 8 | 8 | 8 | CTX, CAZ, CPM, CRO, CFO, NAL, SUT, ATM, TET, CLO |
| KP337 | 0,5 | >32 | >32 | >32 | 4 | 4 | 8 | CTX, CAZ, CPM, CRO, CFO, NAL, ATM, TET, CLO |
| KP326 | 0,5 | >32 | 8 | 8 | 8 | 8 | 16 | CTX, CAZ, CPM, CRO, NAL, SUT, ATM, TET, CLO, GEN |
| KP515 | 0,5 | >32 | >32 | >32 | 8 | 8 | 16 | CTX, CAZ, CPM, CRO, CFO, NAL, SUT, ATM, TET, CLO, FOS |
| KP870 | 0,5 | >32 | >32 | >32 | 16 | 8 | 16 | CTX, CAZ, CPM, CRO, CFO, NAL, SUT, ATM, TET, CLO, FOS |
| FA64 | 1 | 1 | 0.5 | 0.5 | 4 | 8 | 4 | CTX, CAZ, CPM, CRO, NAL, SUT, ATM |
| 2KP | 1 | 8 | 8 | 8 | 8 | 8 | 8 | CTX, CAZ, CPM, CRO, CFO, NAL, SUT, ATM, TET |
| 1ECKPC | 1 | 4 | 8 | 8 | 8 | 8 | 8 | CTX, CAZ, CPM, CRO, CFO, NAL, SUT, ATM, TET, CLO, GEN |
| N7 | 1 | 8 | 8 | 8 | 4 | 8 | 8 | CTX, CAZ, CPM, CRO, NAL, SUT, ATM, TET, CLO, GEN, AMI |

\*POL – polymyxin B, ETP – ertapenem, IMP – imipenem, MER- meropenem, ENO – enrofloxacin, CIP – ciprofloxacin, LVX – levofloxacin, CTX – cefotaxime, CAZ – ceftazidime, CPM – cefepime, CRO – ceftriaxone, CFO – ceftiofur, NAL – nalidixic acid, AMI - amikacin, GEN – gentamicin, SUT – sulfamethoxazole/trimethoprim, ATM - aztreonam, TET – Tetracycline, CLO – chloramphenicol, FOS – fosfomicin.

**Fig. 1.**
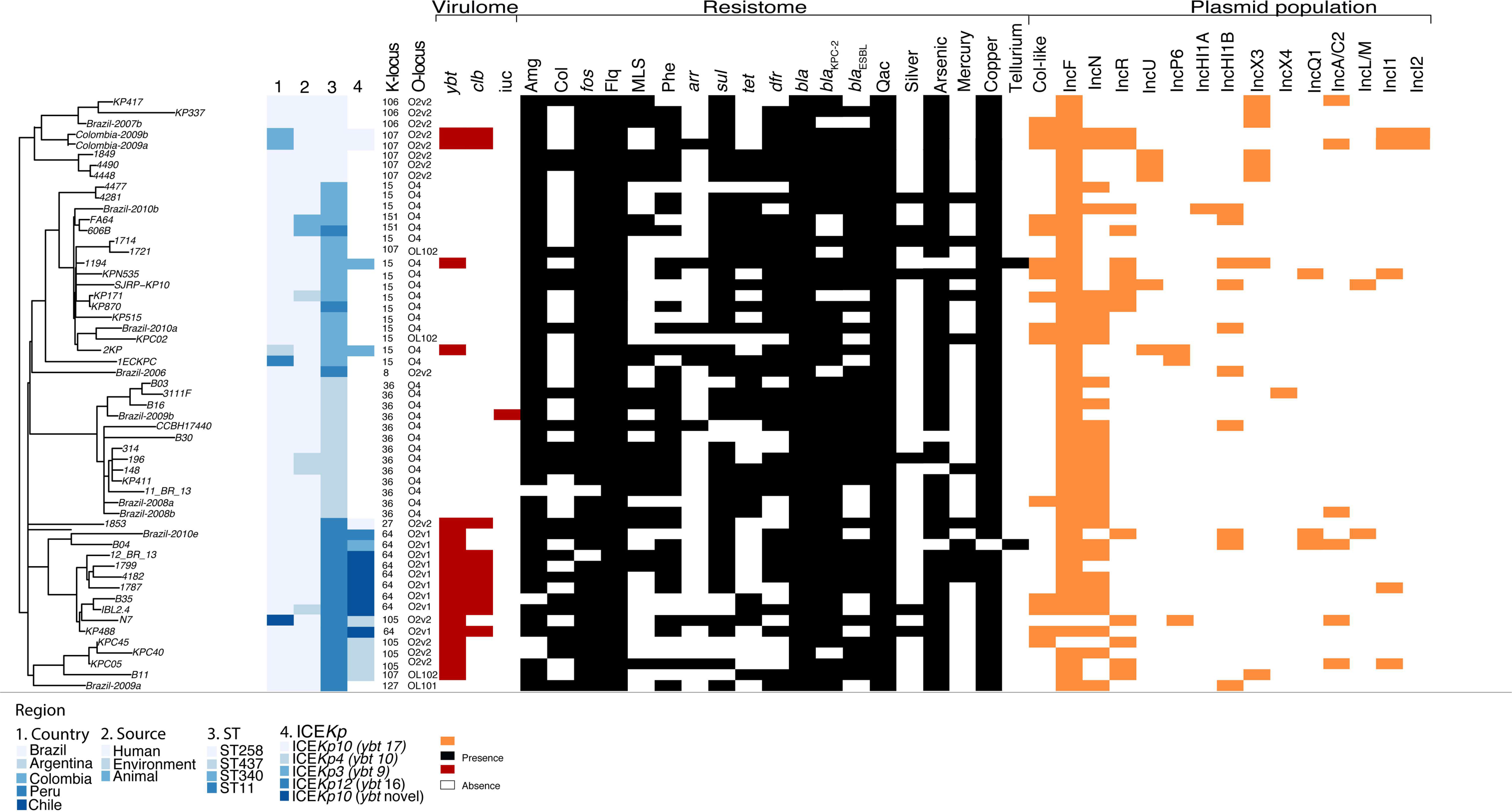
Virulome, resistome and plasmid population of KPC- and/or CTX-M-producing *K. pneumoniae* belonging to CG258 in Latin America. Tracks indicate: (1) country, (2) strain source, (3) sequence type (ST), (4) ICE*Kp* structure. The red/black/orange regions represent the presence of the gene, and blank regions represent their absence. Amg, aminoglycosides resistance genes (i.e., transferases and 16S rRNA methylases); Col, polymyxin resistance genes (including *mgrB*/*pmrB* mutations and *mcr-1*); Flq, fluoroquinolone resistance genes (i.e., QRDR mutations and PMQR); MLS, macrolides resistance genes (*mphA*, *erm*); Phe, phenicols resistance genes (*cat*, *cml*, *flor*) (Table S2).

Additional *in silico* analysis of 36 genome sequences obtained from GenBank confirmed the wider resistome of *K. pneumoniae* strains circulating in Latin America. However, others CTX-M gene variants, such as *bla*_CTX-M-2_, *bla*_CTX-M-8_, *bla*_CTX-M-9_, *bla*_CTX-M-14_ and *bla*_CTX-M-59_ could be identified, as well as the *bla*_KPC-3_ carbapenemase gene identified in two Colombian *K. pneumoniae* strains (Colombia-2009a, Colombia-2009b) alone (Table S2). Moreover, presence of *armA*, *rmtB*, *rmtD* and *rmtG*16S rRNA methyltransferases encoding genes, conferring resistance to most aminoglycosides, was confirmed in 6 of the 55 (11%) analyzed genomes. Worryingly, a third of isolates were predicted to be polymyxin resistant based on deletions in *mgrB* and/or *pmrB* genes in 20 of the 55 analyzed genomes; whereas the *mcr-1* gene was identified in a human strain isolated in Brazil (Table S2).

Plasmid incompatibility (Inc) typing revealed the presence of a wide diversity of plasmid replicon types harbored by KPC-2- and/or CTX-M-type-producing *K. pneumoniae* (Figure 1, Table S2). Among 55 genomes analyzed, IncF, IncN, IncR and Col-like replicons were over-represented (96.4%, 54.5%, 27.3% and 27.3% respectively), whereas other Inc types identified were: X3 (*n*= 8), HI1B (*n*= 8), L/M (*n*= 6), U (*n* = 5), Q1 (*n*= 4), P6 (*n*= 3), A/C2 (*n*= 2), I1 (*n*= 1), I2 (*n*= 1), HI1A (*n*= 1) and X4 (*n*= 1). However, *bla*_KPC_ plasmids belonging to the IncN incompatibility group were identified in 30 of the 55 (54%) carbapenem-resistant *K. pneumoniae* strains.

### Virulome, yersiniabactin, colibactin and integrative conjugative elements (ICEs)

Among genomes analysed, lineages belonging to ST11 and ST340 carried multiple variants of distinct yersiniabactin siderophore (*ybt*) and/or genotoxic colibactin (*clb*) genes from distinct *ybt/clb* lineages and ICE*Kp* variants. In this regard, we detected *ybt* lineage 9 (ICE*Kp3*), *ybt* 10 (ICE*Kp3*) and *ybt* 16 (ICE*Kp3*) in nine *ybt*+ *K. pneumoniae* strains belonging to ST11 and ST340, isolated from human samples collected in Argentina, Brazil and Chile (Figure 2A, Table S3); whereas *ybt* 17 and a novel *ybt* sequence type YbST346, that does not belong to any of the 17 previously described *ybt* lineages (Lam et al, 2018a), were identified together with *clb* lineage 2B in ICE*Kp10* structures, in eight Brazilian *K. pneumoniae* ST11 strains recovered from human and environmental samples (Figure 2B). ST258 lineages from Colombia harbored the classical ICE*Kp10* with the *ybt* 17 and *clb* 2B.

**Fig. 2.**
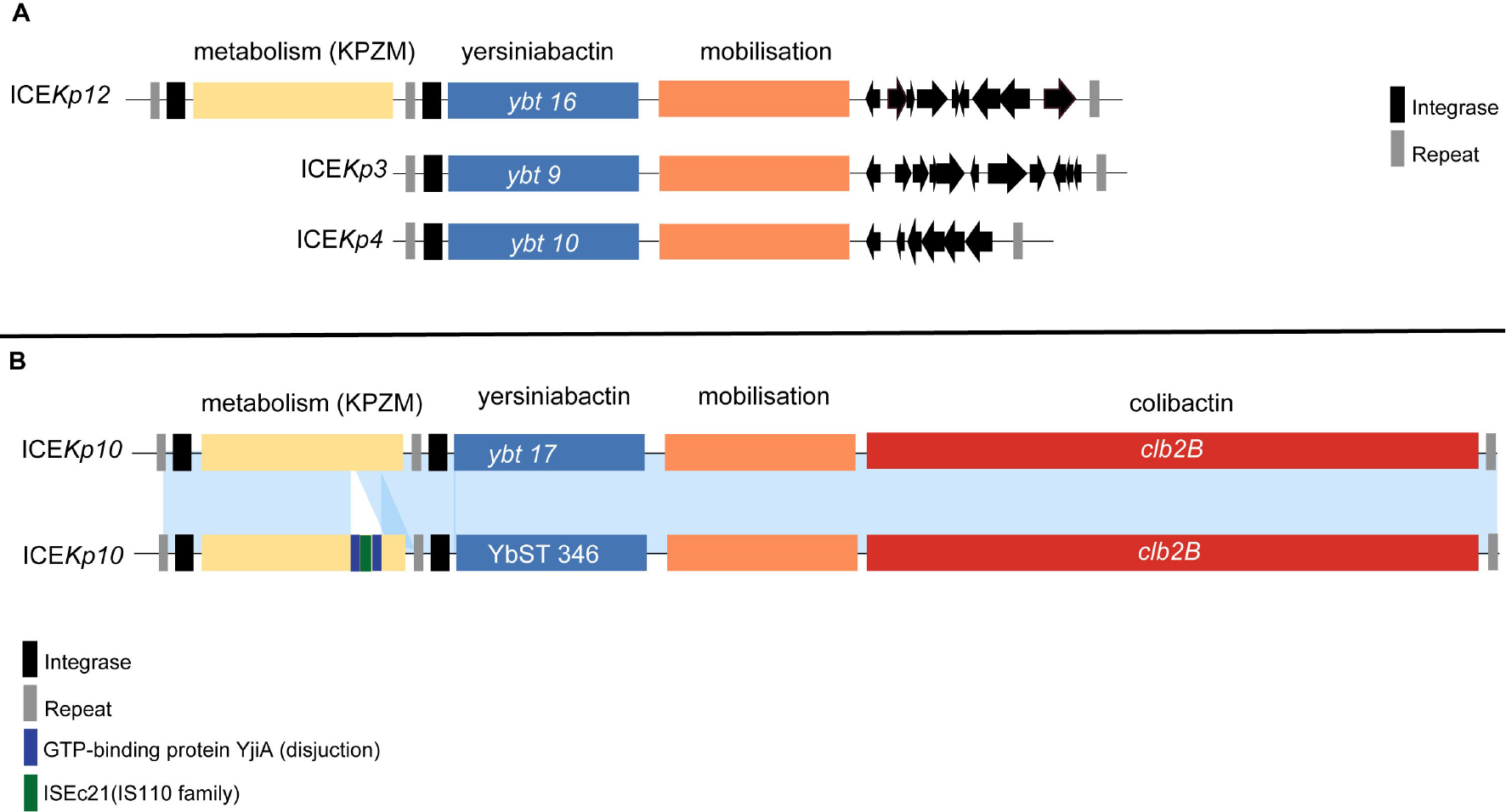
Comparative analysis of integrative and conjugative elements ICE*Kp3*, ICE*Kp4*, ICE*Kp10*, and ICE*Kp12*. In A, ICE*Kp* mobilizing yersiniabactin identified in *K. pneumoniae* CG258 in South America. In B, alignment of ICE*Kp10*/*ybt* 17 against ICE*Kp10* carrying a novel yersiniabactin sequence YbST346, identified in lineages belonging to ST11, isolated in Brazil. Blue blocks represent yersiniabactin synthesis locus *ybt*, labelled with the associated *ybt* lineage. Orange represents the mobilization module. Light orange represents KPZM (Zn2+/Mn2+) metabolism module. Light blue shading denotes shared regions of homology (>95%), where the main difference between the two ICE*Kp10* is the presence of a novel *ybt* lineage, and the insertion sequence IS*Ec21* (IS*110* family) located inside the Zn2+/Mn2+metabolism module (KPZM). The key differences between ICE*Kp3*, ICE*Kp3* and ICE*Kp3* have been previously defined and are restricted to a single variable region, where ICE*Kp3* was constituted by restriction endonuclease, DUF4917 domain containing protein, ATP/GTP phosphatase, reverse transcriptase, DDE endonuclease, and five hypothetical proteins; whereas ICE*Kp4*was formed by transposase, ABC transporter, type I restriction endonuclease, DNA methyltransferase and hypothetical protein; and ICE*Kp12* contained an additional Zn2+/Mn2+metabolism module (KPZM) (Marcoleta et al. 2016; Lam et al., 2018a).

The alignment between the classical ICE*Kp10*/*ybt* 17 and that of ICE*Kp10* with the novel YbST346 shows that the main difference is the insertion sequence ISEc*21* (IS*110* family) located within the Zn2+/Mn2+ metabolism module (KPZM) (Figure 2B, Table S3).

Regarding other virulence determinants, the presence of the aerobactin locus (*iuc*) was only identified in a human KPC-2-positive *K. pneumoniae* strain ST11 from Brazil (Figure 1).

### *In silico* serotyping, capsule locus (KL) analysis and string test

*In silico* serotyping of 55 genomes analyzed showed a predominance of O4 [K36, K15, K-non-typeable (NT)], O2v2 (K8, K27, K-NT) and O2v1 (K64) serotypes, which were associated with ST340 (O4/K15, O4/K-NT), ST437 (O4/K36), ST11 (O2v2/K8, O2v2/27, O2v2/K-NT, O2v1/K64), and ST258 (O2v2/K-NT) (Figure 1, Table S2). On the other hand, we investigated the diversity of capsule synthesis loci using full locus information extracted from whole genome sequences. These results show that K-loci were diverse in human and environmental *K. pneumoniae* ST11 (i.e., KL-8, KL-27, KL-64, KL-105, KL-107, KL-127), in this region (Figure 1, Table S2). Interestingly, in *K. pneumoniae* belonging to ST340, KL-15 was assigned to human and environmental strains collected in Argentina, Peru and Brazil, respectively; whereas KL-151 was only identified in animal strains from Brazil. On the other hand, human and environmental *K. pneumoniae* ST437 (from Brazil) were typed as KL-36; whereas KL-106 and KL-107 accounted for strains of ST258, in Brazil and Colombia, respectively.

KL-64/ST11 and KL-105/ST11showed a high virulence behavior in the *G. mellonella* model, the latter being identified in clinical samples from Chile and Brazil. In this regard, KL-64 has been previously associated with strains from invasive *K. pneumoniae* infections (Follador et al., 2016).

The genetic structure of the cps synthesis loci across the virulent ST11 (KL-105 and KL-64) and ST340 (KL-15) was distinct from the K-loci from hvKpK1 (KL-1) and K2 (KL-2) (Figure 3). In this concern, for these K-loci, a conserved genetic organization at the 5′ end of the *cps* locus was observed from *galF* to *wzc* genes, whereas *wzc-gnd* and *gnd-ugd* regions were variable. Moreover, while in KL-64 the *gnd-ugd* region is composed of genes involved in GDP-D-mannose synthesis (*manB* and *manC*) and deoxythymidine diphosphate (dTDP)-L-rhamnose synthesis (*rmlA*, *rmlB*, *rmlC* and *rmlD*), in KL-105 the *gnd-ugd* region is only composed of operon *manCB* (Figure 3) (Pan et al., 2015; Wyres et al., 2015; Wyres et al., 2016). Interestingly, dTDP-L-rhamnose is the precursor of L-rhamnose, a saccharide required for the virulence of some pathogenic bacteria, being essential for resistance to serum killing and for colonization (Giraud et al., 2000).

**Fig. 3.**
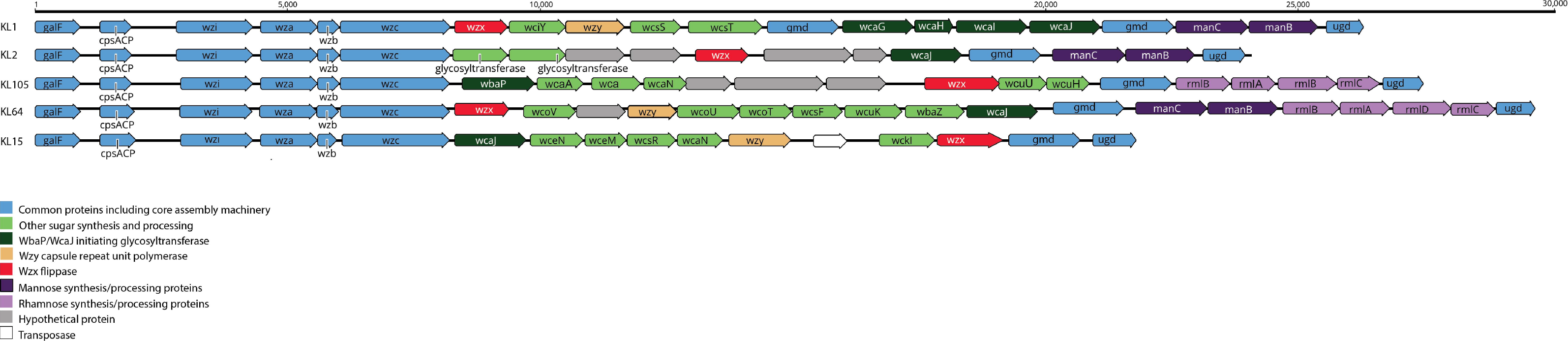
K-loci (KL-1, K-L2, KL-105, KL-64 and KL-15) structures of CR-KP lineages belonging to CG258. In *K. pneumoniae*, K-locus includes a set of genes in the terminal regions encoding for the core capsule biosynthesis machinery (i.e., *galF*, *wzi*, *wza*, *wzb*, *wzc*, *gnd* and *ugd*). The central region is highly variable, encoding for specific sugar synthesis of the capsule, processing and export proteins, plus the core assembly components Wzx (flippase) and Wzy (capsule repeat unit polymerase) (Pan et al., 2015; Wyres et al., 2016). Protein coding sequences are represented as arrows colored by predicted function of the protein product and labelled with gene names where known.

To investigate the hypermucoviscosity phenotype, all the isolates were subjected to the string test. Among the 19 KPC-2 and/or CTX-M-15 producers, only one CTX-M-15-producing *K. pneumoniae* ST340/KL-151 strain (FA64), isolated from a healthy chicken sample, in Brazil, showed hypermucoviscosity. However, neither of the known hypermucoviscosity encoding genes (*rmpA* or *rmpA2*) were detected in its genome.

### *In vivo* virulence behavior of *K. pneumoniae* CG258

Using the *G. mellonella* virulence model, ST11 CR-KP strains (*n*= 2, KL-64/*ybt*+/*clb*+; *n*=2, KL-105/*ybt*+) killed 100% of wax moth larvae inoculated with 1×10^6^ colony-forming units of the bacterial specimens, within 96 h, in a similar way to the known hypermucoviscous hvKp K1/ST23 strain which is *ybt*- and *clb*-negative and carries the pLVPK-like plasmid (*P*> 0.9999) (Figure 4A). KPC-2- and/or CTX-M-15-producing *K. pneumoniae* strains belonging to ST340 killed >60% of wax moth larvae (*n*=2, KL-15/*ybt*+; *n*=3, KL-15/*ybt*ɝ; *n*=1, KL-151/*ybt*-). One ST340 KL-15/*ybt*+ strain isolated from a human infection killed 100% of *G. mellonella* (Figure 4B). *K. pneumoniae* belonging to ST437 (*n*= 3 strains, all KL-36/*ybt*-/*clb*-) killed ~50% of *G. mellonella*, compared to *K. pneumoniae* ATCC 13883 control (Figure 4D).

**Fig. 4.**
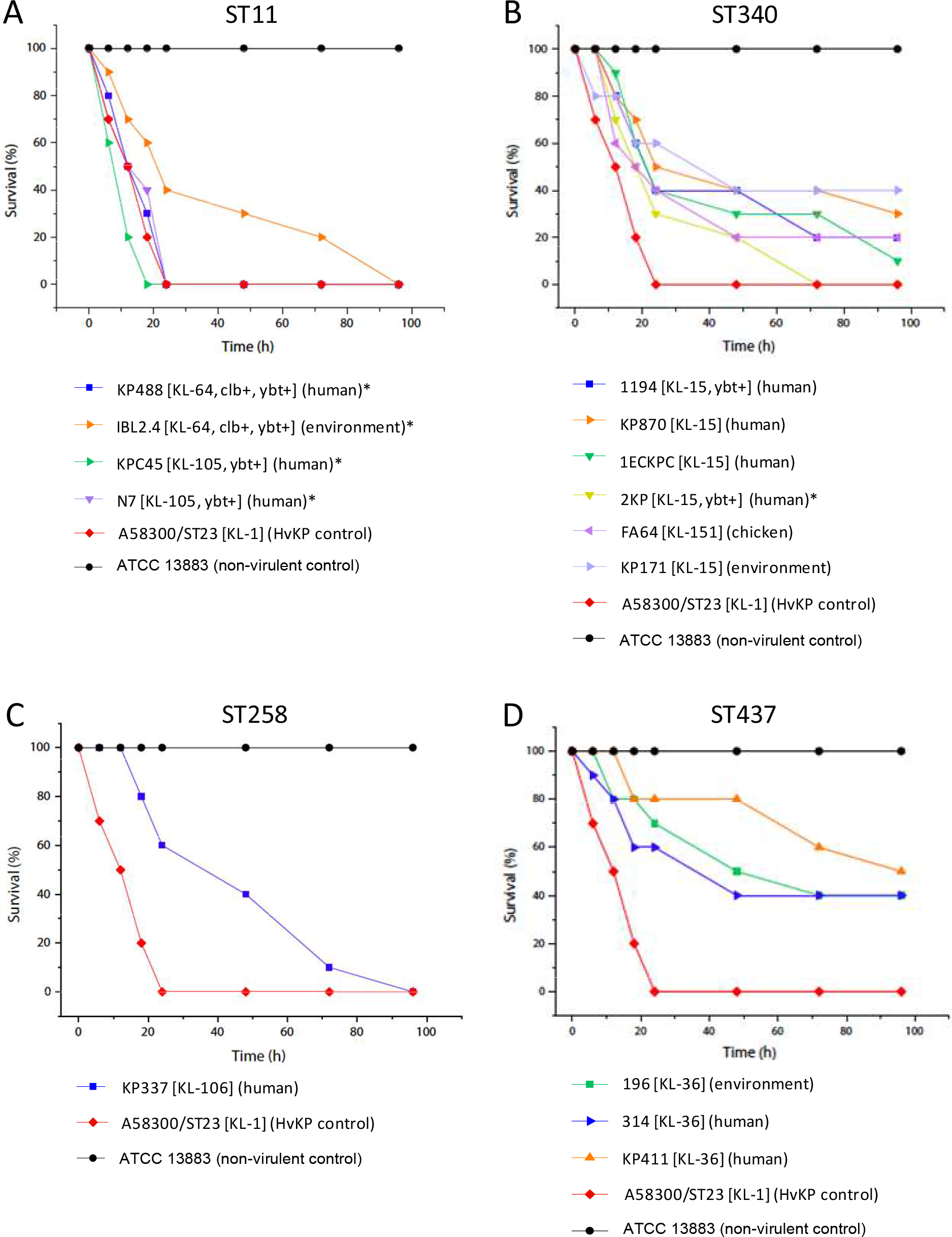
In vivo virulence behavior of CR-KP belonging to ST11, ST340, ST437 and ST258 in a *Galleria mellonella* infection model. The virulence behavior of 1 × 10^6^ colony-forming units of representative *K. pneumoniae* strains on *G. mellonella* survival was assessed using both, non-virulent (ATCC 13883, ST not determined in this study) and hypervirulent (A58300 K1/ST23) *K. pneumoniae* control strains. In A, *K. pneumoniae* KPC45, KP488, IBL2.4 and N7 belonging to ST11, recovered from human and environmental samples. In B, *K. pneumoniae* 1194, KP870, 1ECKPC, 2KP, FA64, KP171 strains belonging to ST340, recovered from human, animal and environmental samples. In C, *K. pneumoniae* KP337 strain belonging to ST258, recovered from a clinical sample. In D, *K. pneumoniae* KP196, KP411 and 314 strains belonging to ST437, recovered from human and environmental samples. Clinical and epidemiological characteristics of *K. pneumoniae* strains are quoted in Table S1. *P>0.9999, indicates no statistically significant difference with respect to the hypervirulent A58300 K1/ST23 *K. pneumoniae* control strain.

Overall, among 14 *K. pneumoniae* strains evaluated, 4/6 (83%) *ybt*+ isolates (4/4 ST11, 1/2 ST340) killed all wax moth larvae within 96 h, compared to only 1/8 (13%) *ybt*-strains (1/1 ST258, 0/4 ST340, 0/3 ST437) (P = 0.03, Fisher’s exact test), suggesting that in the absence of pLVPK-like plasmids, the presence of *ybt* could be enough to confer enhanced virulence. However, the single ST258 strain (*ybt*- and *clb*-negative), isolated from a human clinical sample also killed 100% of *G. mellonella* within 96 h (Figure 4C), suggesting that other factors, not elucidated in this study, may also be contributing to the virulence phenotype of CG258 (Araújo et al., 2018; Hennequin and Robin, 2016; Shah et al., 2017; Fu et al, 2018; Marcoleta et al., 2018; Zheng et al., 2018).

## DISCUSSION

In South American countries, antimicrobial resistance has long been documented to be more challenging than in developed ones (Gales et al., 2012; Sampaio and Gales, 2016). In this regard, the high prevalence of carbapenem resistance in this region has occurred primarily by the dissemination of KPC-producing *K. pneumoniae* isolates belonging to CG258, which have been identified beyond the hospital setting, constituting a One Health problem(Andrade et al., 2011; Gomez et al., 2011; Cejas et al., 2012; Oliveira et al., 2014; Barría-Loaiza et al., 2016; Rojas et al., 2017; Horna et al., 2017; Nascimento et al., 2017). In this study, we performed a resistome and virulome analysis of KPC-and/or CTX-M-producing *K. pneumoniae* lineages belonging to CG258, circulating in hospital settings. Additionally, environmental and animal *K. pneumoniae* isolates, recovered in Brazil, were also investigated.

Genome analysis revealed a wider resistome, which includes genetic determinants conferring resistance to human and animal antibiotics, QACs and HMs, supporting persistence and adaptation of CG258 to different hosts and anthropogenically affected environments. Among MDR and PDR lineages, the presence of mutations in *mgrB*/*pmrB* genes and in the quinolone resistance-determining region; as well as acquisition of 16s rRNA methylases- and β-lactamases-encoding genes (includying *bla*_ESBL_ and *bla*_KPC-2_) have contributed with resistance to polymyxins, fluoroquinolones, aminoglycosides and broad-spectrum β-lactam antibiotics. Moreover, we have identified, for the first time, the presence of the narrow-spectrum β-lactamase encoding gene *bla*_LAP-2_ (GenBank accession number EU159120) and *bla*_TEM-55_ ESBL gene(GenBank accession number DQ286729) in *K. pneumoniae* strains ST340 recovered from swine and human hosts, respectively, in Brazil, confirming versatility of this lineage to acquire novel genetic determinants of resistance.

We have identified regional *bla*_KPC_ spread consistent with high prevalence of IncN plasmids, previously associated with the global spread of these genes (Stoesser et al., 2017). On the other hand, the wide diversity of Inc-type plasmids, found in this study, including small mobilizable Col-like replicons could be associated with the acquisition of multiple resistance mechanisms, contributing to the wider resistome. Therefore, the presence of *K. pneumoniae* in a wide range of environmental reservoirs and hosts, with plasmids that have been shown to facilitate the dissemination of successful resistance genes, even in the absence of selection pressures, may represent a difficult situation to control (Stoesser et al., 2017). Another important issue is the identification of IncHI1-type plasmids, which have been associated with the dissemination of *mcr-1* and *bla*_CTX-M_-type genes in Colombia and Uruguay, respectively (Saavedra et al., 2017; Garcia-Fulgueiras et al., 2017).

Hypervirulent *K. pneumoniae* strains have been sporadically reported in Argentina and Brazil, being associated with remarkable mortality and the production of a hypermucoviscous phenotype in lineages belonging to ST23 with capsular serotype K1, and ST29/K19 (Cejas et al., 2014; Coutinho et al., 2014, Moura et al., 2017). In this study, virulome analysis revealed that *ybt* and *clb* genes have been acquired by strains of CG258 in South America, highlighting the need to also consider these additional virulence factors rather than the presence of pLVPK-like plasmids and hypermucoviscous phenotypes alone, in the establishment of hypervirulence in carbapenem-resistant lineages of *K. pneumoniae*.

Notably, there have been increasing reports of highly virulent *K. pneumoniae* strains carrying *ybt* belonging to the international clone ST11. The emergence of CR-hvKp strains carrying *ybt* plus a deletion variant of the pLVPK-like plasmid belonging to ST11 has been associated with outbreak of fatal nosocomial infection in China (Gu et al., 2018), raising an epidemiological alert in response to the increased number of cases reported, in the last year (Lee et al., 2017; Zhan et al., 2017; Chen and Kreiswirth, 2018; Du et al., 2018; Wong et al., 2018; Yao et al., 2018).

In summary, these results (available for interactive exploration in Microreact at https://microreact.org/project/H1LSZsRz7) confirm the enhanced virulence of KPC-2- and/or CTX-M-producing *K. pneumoniae* belonging to the international high-risk clone CG258 in South America, where acquisition of ICE*Kp* encoding yersiniabactin and colibactin, and wider resistome have likely contributed to enhanced virulence and persistence of ST11 (KL-64 and KL-105) and ST340 (KL-15) lineages, in the human-environment interface. While capsule composition deserves further investigation, active surveillance should not only focus on clonal origin, antimicrobial resistance and presence of pLVPK-like plasmids, but also the virulence associated with yersiniabactin and colibactin, as well as other biomarkers for differentiation of hvKp from classical *K. pneumoniae* (Russo et al., 2018); and control measures should be conducted to prevent the global dissemination of these lineages.

## CONCLUSION

Our study points out several important issues. Firstly, interplay of yersiniabactin and/or colibactin and KPC-2 production has become to be identified among *K. pneumoniae* belonging to CG258, in South America, contributing to the emergence of highly virulent lineages that pose great risk to human health (Lam et al. 2018a). Second, in South America ICE*Kp3*, ICE*Kp3*and ICE*Kp10*carrying *ybt* and/or *clb* circulate among KPC-2-producing*K. pneumoniae* belonging to ST11 (KL-64 andKL-105), where multiple distinct K-loci often indicates distinct sublineages that may correlate with independent ICE*Kp* acquisitions (supported by our phylogenetic analysis shown in Figure 1);being associated with enhanced virulence, these should be considered a target for genomic surveillance along with antimicrobial resistance determinants. Third, the wide resistome could be contributing to adaptation of KPC-2- and/or CTX-M-producing *K. pneumoniae* CG258 in the human-animal-environment interface, highlighting the urgent need for enhanced control efforts. Finally, these findings could contribute to the development of strategies for prevention, diagnosis and treatment of *K. pneumoniae* infections.

## ACKNOWLEDGEMENTS

Fundação de Amparo à Pesquisa do Estado de São Paulo (FAPESP2016/08593-9) and Conselho Nacional de Desenvolvimento Científico (CNPq462042/2014-6) research grants are gratefully acknowledged. L.C. and N.L. are research grant fellows of FAPESP (2015/21325-0) and CNPq (312249/2017-9), respectively. K.E.H. is supported by a Senior Medical Research Fellowship from the Viertel Foundation of Australia. We thank Cefar Diagnóstica Ltda. (Brazil) for kindly supplying antibiotic discs for susceptibility testing, the Australian Genome Research Facility (AGRF) for PromethION sequencing, and Drs Marcia M. Morais, Ana C. Gales, Andrea M. Moreno, Ketrin Silva and Mara Nogueira for kindly provide *K. pneumoniae* strains and/or fastq files.

## Abbreviations

CR-hvKp: carbapenem-resistant hypervirulent *Klebsiella pneumoniae*
KPC: *Klebsiella pneumoniae* carbapenemase
CTX: cefotaximase
CG: clonal group
ST: sequence type
QACs: quaternary ammonium compounds
KL: K-locus
ybt: yersiniabactin
clb: colibactin
ICEKp: integrative conjugative element *K. pneumoniae*
pLVPK: large virulence plasmid of *K. pneumoniae*
CPS: capsular polysaccharides
MLST: multilocus sequence typing
YbSTs: yersiniabactin sequence types
CbSTs: colibactin sequence types
CR-Kp: carbapenem-resistant *K. pneumoniae*
MIC: minimum inhibitory concentration
ESBL: extended-spectrum beta-lactamase
HM: heavy metal
ML: maximum likelihood
MDR: multidrug resistance
PDR: pandrug resistance
Inc: incompatibility
IS: insertion sequence
KPZM: Zn2+/Mn2+metabolism module
QRDR: quinolone-resistance determining region
PMQR: plasmid-mediated quinolone resistance.

